# Glucoregulation and coping behavior after chronic stress: sex differences across the lifespan

**DOI:** 10.1101/2021.03.17.435842

**Authors:** Carley Dearing, Rachel Morano, Elaine Ptaskiewicz, Parinaz Mahbod, Jessie R Scheimann, Ana Franco-Villanueva, Lawson Wulsin, Brent Myers

**Affiliations:** Biomedical Sciences, Colorado State University, Fort Collins, CO; Pharmacology and Systems Physiology, University of Cincinnati, Cincinnati, OH; Psychiatry and Behavioral Neuroscience, University of Cincinnati, Cincinnati, OH

**Keywords:** forced swim test, corticosterone, glucose tolerance, adolescence, aging

## Abstract

Exposure to prolonged stress during adolescence taxes adaptive and homeostatic processes leading to deleterious behavioral and metabolic outcomes. Although previous pre-clinical studies found effects of early life stress on cognition and stress hormone reactivity, these studies largely focused on males. The purpose of the current study was to determine how biological sex shapes behavioral coping and metabolic health across the lifespan after chronic stress. We hypothesized that examining chronic stress-induced behavioral and endocrine outcomes would reveal sex differences in the biological basis of susceptibility. During the late adolescent period, male and female Sprague-Dawley rats experienced chronic variable stress (CVS). Following completion of CVS, all rats experienced a forced swim test (FST) followed 3 days later by a fasted glucose tolerance test (GTT). The FST was used to determine coping in response to a stressor. Endocrine metabolic function was evaluated in the GTT by measuring glucose and corticosterone, the primary rodent glucocorticoid. Animals then aged to 15 months when the FST and GTT were repeated. In young animals, chronically stressed females exhibited more passive coping and corticosterone release in the FST. Additionally, chronically stressed females had elevated corticosterone and impaired glucose clearance in the GTT. Aging affected all measurements as behavioral and endocrine outcomes were sex specific. Furthermore, regression analysis between hormonal and behavioral responses identified associations depending on sex and stress. Collectively, these data indicate female susceptibility to the effects of chronic stress during adolescence. Further, translational investigation of coping style and glucose homeostasis may identify biomarkers for stress-related disorders.

## Introduction

The neuroendocrine stress response, initiated by threats to homeostasis (Chrousos and Gold, 1992), is essential for mobilizing energy necessary for adaptation and survival (Myers et al., 2014). However, dysregulated or prolonged stress responses are linked to cardiometabolic dysfunction and neuropsychiatric disorders (Sudheimer et al., 2015). Globally, metabolic dysfunction contributes to leading causes of death such as cardiovascular disease (Anand et al., 2003) and diabetes mellitus (Isfort et al., 2014). Additionally, metabolic and depressive disorders have a bi-directional relationship, further contributing to years lived with disability (McIntyre et al., 2009; Petrie et al., 2018).

While the development of metabolic dysfunction is multifactorial, stress is widely recognized as a causative agent (Picard et al., 2014). Central stress responses influence the neuroendocrine hypothalamic-pituitary-adrenal (HPA) axis, as well as the sympathetic-adrenal-medullary (SAM) axis. Thus, stressful events initiate a cascade of neural and endocrine responses that mobilize energy to alter both physiologic and psychologic states (Kyrou and Tsigos, 2009). Aberrant activation of these systems by prolonged early life stress is linked to the development of metabolic dysfunction in adulthood (Kaufman et al., 2007; Vargas et al., 2016), as well as psychiatric conditions, including depressive disorders (Carr et al., 2013; Enoch, 2011). In rodent models, chronic early life stress increases HPA axis reactivity (Jankord et al., 2011; Vargas et al., 2016) and impairs neuronal survival and cognition in adulthood (Naninck et al., 2015). Additionally, chronic variable stress (CVS) in young adult rats increases fasting insulin and corticosterone in males (Pereira et al., 2016).

Although pre-clinical studies have found immediate and long-term effects of early life stress on cognitive function (Bolton et al., 2017; Brunson et al., 2005; Hedges and Woon, 2011; Naninck et al., 2015; Pechtel and Pizzagalli, 2011; Saleh et al., 2017) and endocrine stress reactivity (Goldman-Mellor et al., 2012; Loman and Gunnar, 2010; McEwen, 2009; Vargas et al., 2016), these studies have largely focused on males.

While there are significant sex differences in the development of psychiatric disorders (Jalnapurkar et al., 2018; Oliveira et al., 2017), we are just beginning to learn how biological sex impacts behavioral and metabolic health. Further, studies examining potential sex differences in coping and HPA axis responses after stress in adolescence have generated equivocal results (Cotella et al., 2020; Wulsin et al., 2016), possibly related to group variation and/or timing of assessments.

In the current study, we employed a longitudinal design to test the hypothesis that females may have differential behavioral and metabolic susceptibility to adolescent chronic stress. Furthermore, we sought to examine how early life measures may predict an individual’s behavioral and endocrine outcomes later in life. To this end, we assessed the effects of chronic stress during adolescence on coping behavior and glucoregulation in a large cohort of male and female rats both immediately following stress exposure and at 15 months of age. This approach permitted the analysis of stress, sex, and age interactions on coping behaviors, as well as glucocorticoid and glucose responses to psychological and physiological challenges.

## Methods

### Animals

Animals were kept on a 12 on 12 off light-dark cycle. Males were kept on a 6:00-18:00 cycle and females were kept on a 9:00-21:00 cycle to maintain the same circadian time during experimentation. Food and water were available *ad libitum*. All experiments were approved by the Institutional Animal Care and Use Committee of the University of Cincinnati (protocol 04-08-03-01) and complied with the National Institutes of Health Guidelines for the Care and Use of Laboratory Animals. All animals had daily welfare assessments by veterinary and/or animal medical service staff.

### Design

As illustrated in **Fig. 1**, a single cohort of rats was generated through in-house breeding of 12 litters of 10 Sprague-Dawley rats each to produce 120 pups (60 rats/sex). Pups were cross fostered as necessary to maintain uniform litter size and equal sex distribution (5 pups/sex/litter). All pups were weaned at postnatal (PN) day 24. At 5 weeks of age, pups were separated into treatment-specific same sex pairs so that 3 rats/sex/liter were assigned to the CVS group and 2 rats/sex/litter were assigned to No CVS. Pairings were randomized and not limited to littermates. In total, 36 rats/sex were exposed to CVS and 24 rats/sex remained as unstressed controls. All rats were handled regularly with body weight monitored throughout the course of the experiment. Males and females were housed separately in adjacent rooms with light cycles shifted so that all stressors and assessments occurred at the same time each day for both sexes. Due to the scale of the experiment, equal portions of all 4 groups went through sample collection and testing on consecutive days. CVS began at PN 43 with basal blood samples taken 14 days later. After 20 days of CVS, rats went through the forced swim test (FST) followed 3 days later by a fasted glucose tolerance test (GTT). Animals then aged for 13 months prior to another basal blood sample, FST, and GTT. Tissues were then collected for analysis 3 days later at just over 15 months of age.

**Figure 1:**
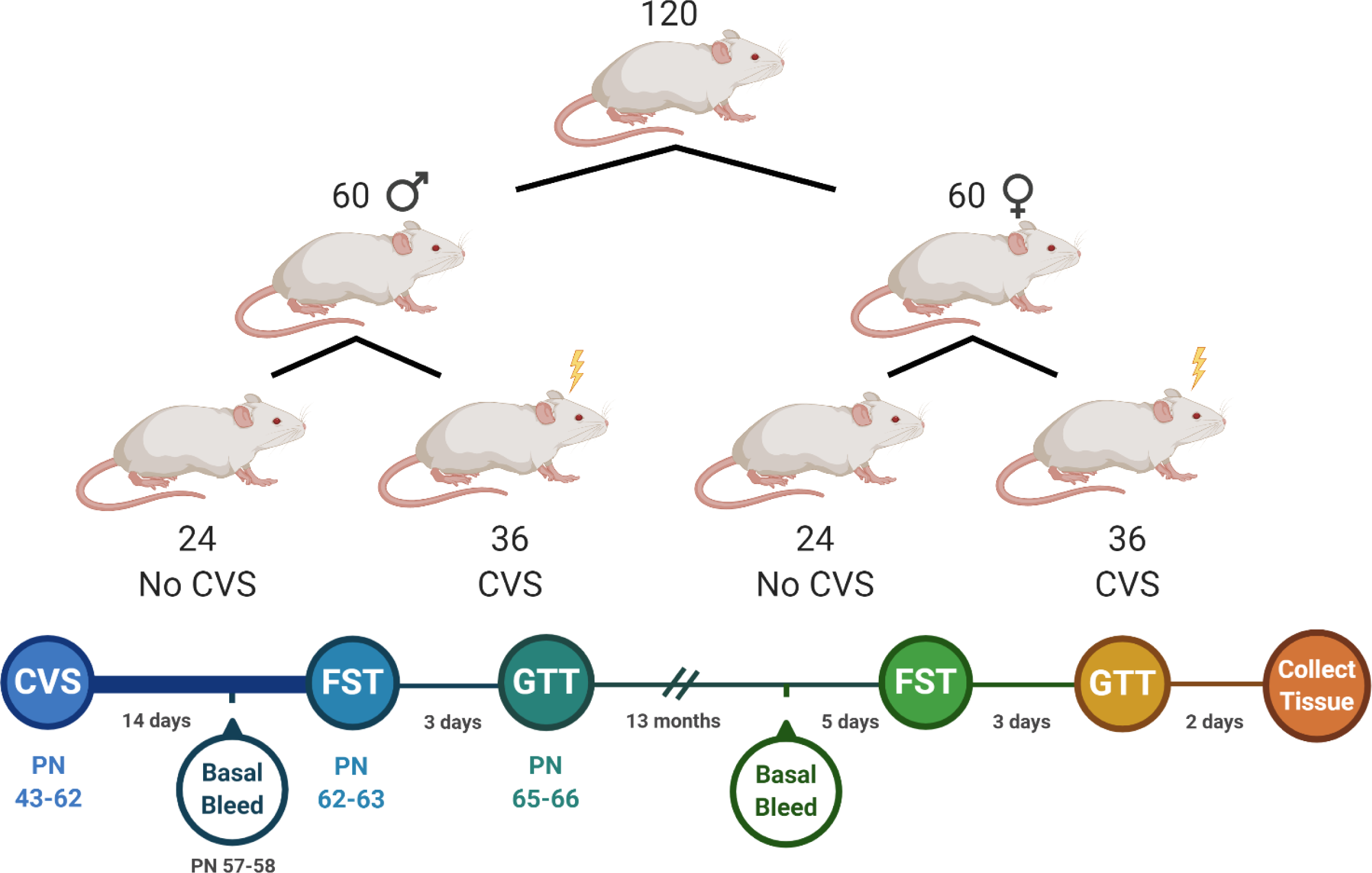
Experimental design and timeline. A single cohort of rats was separated into 4 experimental groups consisting of male and female unstressed controls (No CVS, n = 24/sex) and chronically stressed males and females (CVS, n = 36/sex). These groups underwent endocrine and behavioral assessments immediately following CVS. After aging for 13 months, they were re-tested prior to tissue collection.

### CVS

Exposure to CVS began at PN 43, which is considered late adolescence (Jankord et al., 2011). CVS continued for 20 days to include the transition to adulthood. Animals were exposed to stressors twice daily (AM and PM) presented in a randomized manner (Jankord et al., 2011; Myers et al., 2017; Schaeuble et al., 2019). Stressors included restraint (plexiglass tube, 30 min), shaker (100 RPM, 1 hr), damp bedding (1 hr), cold room (4 °C, 1 hr), and hypoxia (8% oxygen, 30 min). Animals were also exposed to overnight stressors twice a week. These included housing with an unfamiliar cage mate and single housing for social instability and social isolation, respectively.

### Blood collection and analysis

Blood samples (approximately 100 µL) were collected by tail clip in tubes containing 10 µL of 100 mmol/L ethylenediamine tetraacetate (Myers et al., 2017). Baseline blood was collected on the morning of day 15 in the CVS paradigm prior to daily stressors. Following CVS, animals were exposed to novel acute stressors that acted as psychologic (FST) and glycemic (GTT) challenges. All 3 assessments were repeated 13 months later. Blood glucose was determined from tail blood using Bayer

Contour Next glucometers and test strips (Ascensia, Parsippany, NJ). Collected blood samples were centrifuged at 3000× g for 15 minutes at 4°C and plasma was stored at - 20°C until analysis. Triglyceride and cholesterol levels were determined by the Mouse Metabolic Phenotyping Center at the University of Cincinnati. Plasma corticosterone was measured using an ENZO Corticosterone ELISA (ENZO Life Sciences, Farmingdale, NY) with an intra-assay coefficient of variation of 8.4% and an inter-assay coefficient of variation of 8.2% (Bekhbat et al., 2018).

### FST

The FST was used as psychological stressor to assess active versus passive coping (Molendijk and de Kloet, 2019) and endocrine reactivity. As previously described (Pace et al., 2020), rats were placed in an open-top cylinder (61 cm high x 19 cm diameter) filled with 40 cm of water (23-27 °C, 10 minutes). Behavior was recorded with an overhead mounted camera and scored by a blinded observer. The video was analyzed every 5 s for observations of immobility (passive coping) or activity (swimming, climbing, and diving). Additionally, blood was collected at 15, 30, 60, and 120 minutes after the beginning of FST. For females, vaginal swabs were taken after each FST and analyzed for estrous phase by blinded observers.

### GTT

Three days after the FST, animals were fasted for 4 hours prior to a GTT to assess glucose homeostasis and glucocorticoid responses to glycemic challenge (Ghosal et al., 2015; Smith et al., 2014). A baseline blood sample was collected prior to glucose injection (1.5 g/kg, 20% glucose, i.p.). Blood was then collected at 15, 30, 45, and 120 minutes post injection. For females, vaginal swabs were taken after each GTT and analyzed for estrous phase by blinded observers.

### Tissue collection

Two days after the final GTT, animals were rapidly anesthetized (5% inhaled isoflurane) and euthanized via rapid decapitation. Mesenteric white adipose tissue (mWAT), inguinal white adipose tissue (iWAT), and adrenal glands were collected and weighed. mWAT was collected to sample visceral adiposity and iWAT to sample subcutaneous adiposity (Hill et al., 2020; Nguyen et al., 2019). All raw organ weights were corrected for bodyweight.

### Data analysis

Data are expressed as mean ± standard error of the mean. All animals were included in all analyses. All data were analyzed using Prism 8 (GraphPad, San Diego, CA), with statistical significance set at p < 0.05 for all tests. Body weight during CVS, as well as plasma corticosterone and blood glucose during FST and GTT were analyzed by 3-way mixed effects analysis with sex, stress condition, and time (repeated) as factors. In the case of significant main or interaction effects, Tukey multiple comparison post-hoc tests were used to compare groups at specific times. Basal plasma measures, FST behavior, hormonal area under the curve (AUC), aged body weight, and organ weights were analyzed with 2-way ANOVA with sex and stress condition as factors. For main or interaction effects, Tukey multiple comparison post-hoc tests were employed.

Pearson correlation two-tailed analyses were run to examine associations between young and aged data within each group.

## Results

### Body weight and basal plasma measures during CVS

Body weight analysis over the course of CVS (**Fig. 2A**) found main effects of sex [F(1, 116) = 480.5, p < 0.0001], CVS [F(1, 116) = 6.866, p = 0.01], and day [F(1.590, 184.5) = 3216, p < 0.0001], as well as sex x day [F(5, 580) = 341.7, p < 0.0001] and CVS x day [F(5, 580) = 4.439, p < 0.0006] interactions. Post-hoc testing indicated that males had greater body weight than females within stress conditions on all days.

**Figure 2:**
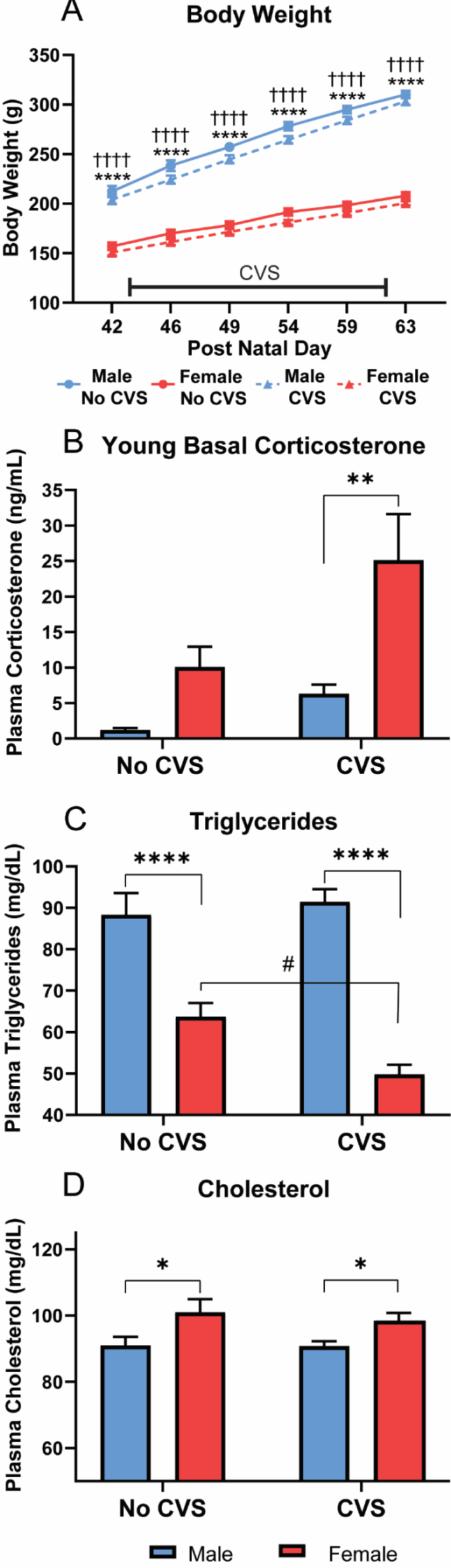
Young metabolic measures. Body weight was measured throughout the course of CVS for both CVS (n = 36/sex) and No CVS (n = 24/sex) controls (**A**). On the morning of CVS day 14, a basal blood sample was taken to determine plasma corticosterone (**B**), plasma triglycerides (**C**), and plasma cholesterol levels (**D**). Data are expressed as mean ± SEM. * represents sex differences within stress condition, ^#^ represents CVS effects within sex, ^†^ represents sex differences within CVS over time. *,^#^ p < 0.05, ** p < 0.01, and ****,^††††^ p < 0.0001.

Although CVS x day interactions were present across sexes, no significant effects of CVS were observed within sex on specific days. Basal corticosterone (**Fig. 2B**) had main effects of sex [F(1, 113) = 10.19, p = 0.0018] and CVS [F(1, 113) = 5.410, p = 0.0218]. Post-hoc analysis found that, within CVS, females had greater (p < 0.01) corticosterone than males. Although, the effect of CVS within females was not significant (p < 0.07). Triglycerides (**Fig. 2C**) showed effects of sex [F(1, 114) = 92.19, p < 0.0001] and sex x CVS interaction [F(1, 114) = 6.161, p = 0.0145]. Specifically, males had higher triglycerides than females (p < 0.0001) in both stress conditions and CVS decreased (p < 0.05) triglycerides in females. For plasma cholesterol (**Fig. 2D**), there was a main effect of sex [F(1, 114) = 17.61, p < 0.0001] where females had elevated cholesterol (p < 0.05) compared to males in both stress conditions. Taken together, these data indicate stress-independent sex differences in triglyceride and cholesterol regulation while CVS increases corticosterone and lowers triglycerides in females.

### Young FST

Behavior during the FST was quantified to determine coping responses to acute psychological stress. Immobility (**Fig. 3A**), a passive behavioral response, showed effects of sex [F(1, 116) = 32.06, p < 0.0001] and CVS [F(1, 116) = 3.944, p = 0.0494] where females were more immobile (p < 0.01) than males in both stress conditions. Although there was a main effect of CVS, within sex there were no significant differences in immobility (males p = 0.061). No significant sex or stress effects were found for climbing or diving behaviors; however, swimming (**Fig. 3B**) showed both sex [F(1, 116) = 16.62, p < 0.0001] and CVS [F(1, 116) = 8.895, p = 0.0035] effects. Post-hoc analysis found that, in the No CVS groups, females had less (p < 0.01) swimming behavior. CVS decreased swimming in males (p < 0.05) with no significant difference (p = 0.078) from females.

**Figure 3:**
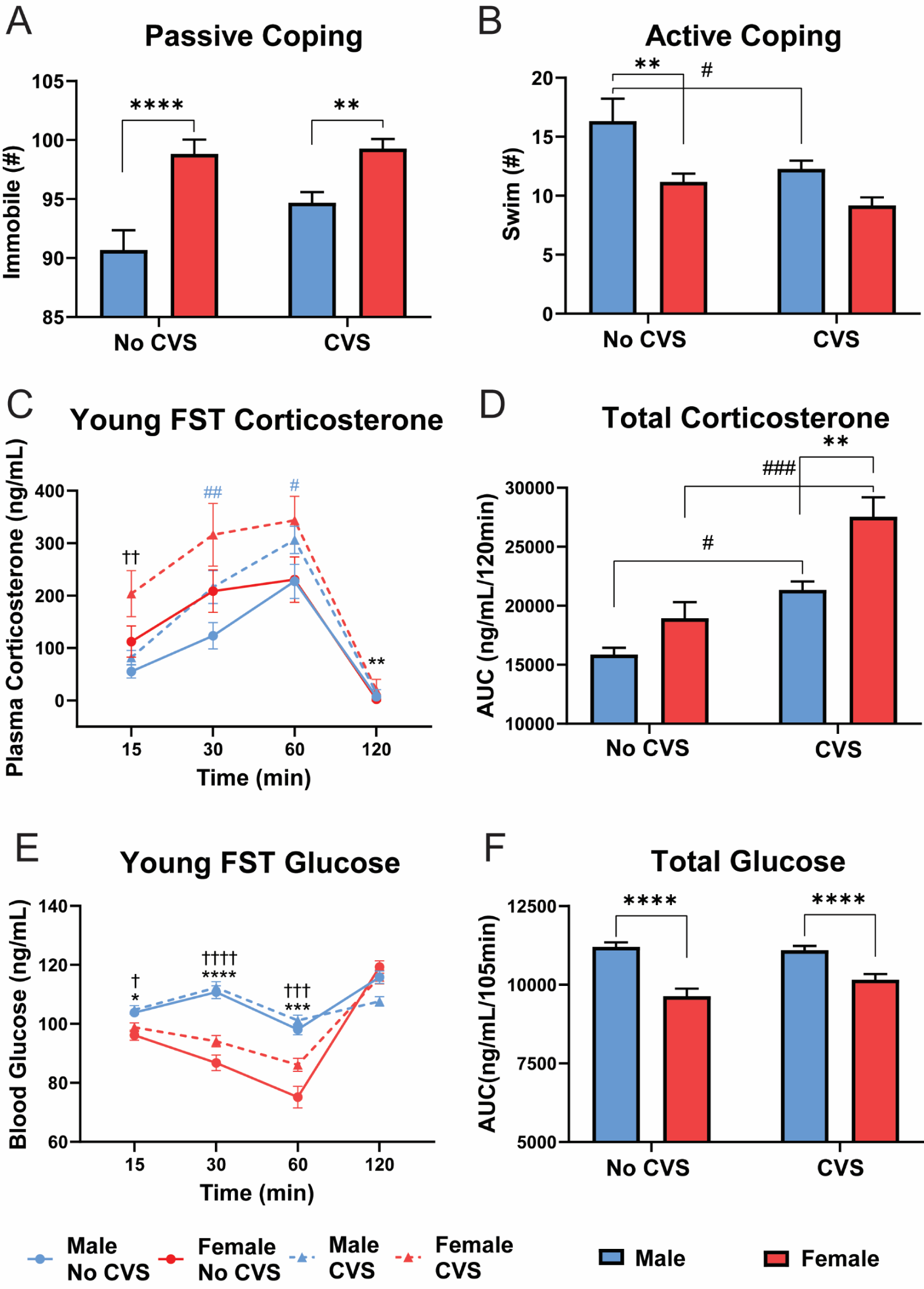
Young FST. During FST, behavioral coping was assessed as passive (**A**) or active (**B**) in No CVS (n = 24/sex) and CVS (n = 36/sex) rats. Blood was taken at 15, 30, 60, and 120 min after the initiation of the 10-min FST. Plasma corticosterone was measured (**C**) and the integrated total corticosterone response was calculated from the AUC (**D**). Blood glucose was also measured at each time point (**E**) and total blood glucose calculated (**F**). Data are expressed as mean ± SEM. * represents sex differences within stress condition, ^#^ represents CVS effects within sex (indicated by color for time-dependent measures), ^†^ represents sex differences within CVS over time. *,^#^,^†^ p < 0.05, **,^##^,^††^ p < 0.01, ***,^###^,^†††^ p< 0.001, and ****,^††††^ p < 0.0001.

Corticosterone (**Fig. 3C**) was measured to determine sex and stress effects on HPA axis reactivity to an acute behavioral challenge. Although all other data are represented by n = 24/sex No CVS and n = 36/sex CVS, a shipping error resulted in lost blood samples for the young FST corticosterone measurement (1/4 randomly across all groups and timepoints). Consequently, the reduced sample sizes are n = 10-24 male No CVS, n = 12-24 female No CVS, n = 17-36 male CVS, and n = 18-36 female CVS. Mixed effects analysis found main effects of sex [F(1, 116) = 32.16, p < 0.0001], CVS [F(1, 116) = 55.03, p < 0.0001], and time [F(2.413, 174.3) = 462.2, p < 0.0001] with sex x time [F(4, 289) = 11.79, p < 0.0001] and CVS x time [F(4, 289) = 13.77, p < 0.0001] interactions. Post-hoc analysis indicated CVS females had higher corticosterone than CVS males 15 min post-FST, while CVS increased corticosterone (p < 0.05) in males at 30 and 60 min. Additionally, females had higher corticosterone (p < 0.01) at the 90 min recovery time point. AUC analysis of cumulative corticosterone exposure during the FST (**Fig. 3D**) found effects of sex [F(1, 48) = 11.95, p = 0.0012] and CVS [F(1,48) = 27.49, p < 0.0001]. Specifically, CVS increased total corticosterone (p < 0.05) in both sexes but to a greater extent (p < 0.01) in females than males.

Glucose mobilization is a component of the stress response reflecting metabolic integration of sympathetic-mediated glucagon release, gluconeogenesis, and glycogenolysis, in addition to longer-term glucocorticoid effects on gluconeogenesis and glycolysis (Lin et al., 2021; Tank and Lee Wong, 2015). FST exposure (**Fig. 3E**) led to effects of sex [F(1, 116) = 44.30, p < 0.0001] and time [F(2.47, 284.4) = 151.8, p < 0.0001], with sex x time [F(3, 346) = 59.55, p < 0.0001] and CVS x time [F(3, 346) = 10.74, p < 0.0001] interactions. From 15-60 min, females had lower (p < 0.05) blood glucose than males regardless of stress condition. This was reflected by significant sex effects [F(1,114) = 48.94, p < 0.0001] in the AUC analysis (**Fig. 3F**). Collectively, the young FST data indicate that, compared to males, females more passively cope with behavioral challenge and have greater CVS-induced glucocorticoid hypersecretion.

### Young GTT

The GTT was utilized to examine sex and CVS effects on glucose tolerance, as well as glucocorticoid responses to a systemic hyperglycemia stressor. Blood glucose responses to i.p. glucose (**Fig. 4A**) had effects of time [F(2.41, 279.1) = 387.8, p < 0.0001], sex x CVS [F(1, 116) = 5.15, p = 0.025], sex x time [F(4, 464) = 12.10, p < 0.0001], and sex x CVS x time [F(4, 464) = 2.796, p = 0.0257] interactions. Post-hoc analysis found that peak glucose (15 min) was greater (p <0.001) in CVS females than CVS males. Further, CVS males had enhanced glucose clearance (p <0.01) 45 min post-injection. Cumulative glucose exposure (**Fig. 4B**) also showed sex x CVS interaction [F(1, 116) = 4.703, p = 0.0322] where CVS decreased total glucose in males (p < 0.05) and, within CVS, females had greater (p < 0.05) total glucose.

**Figure 4:**
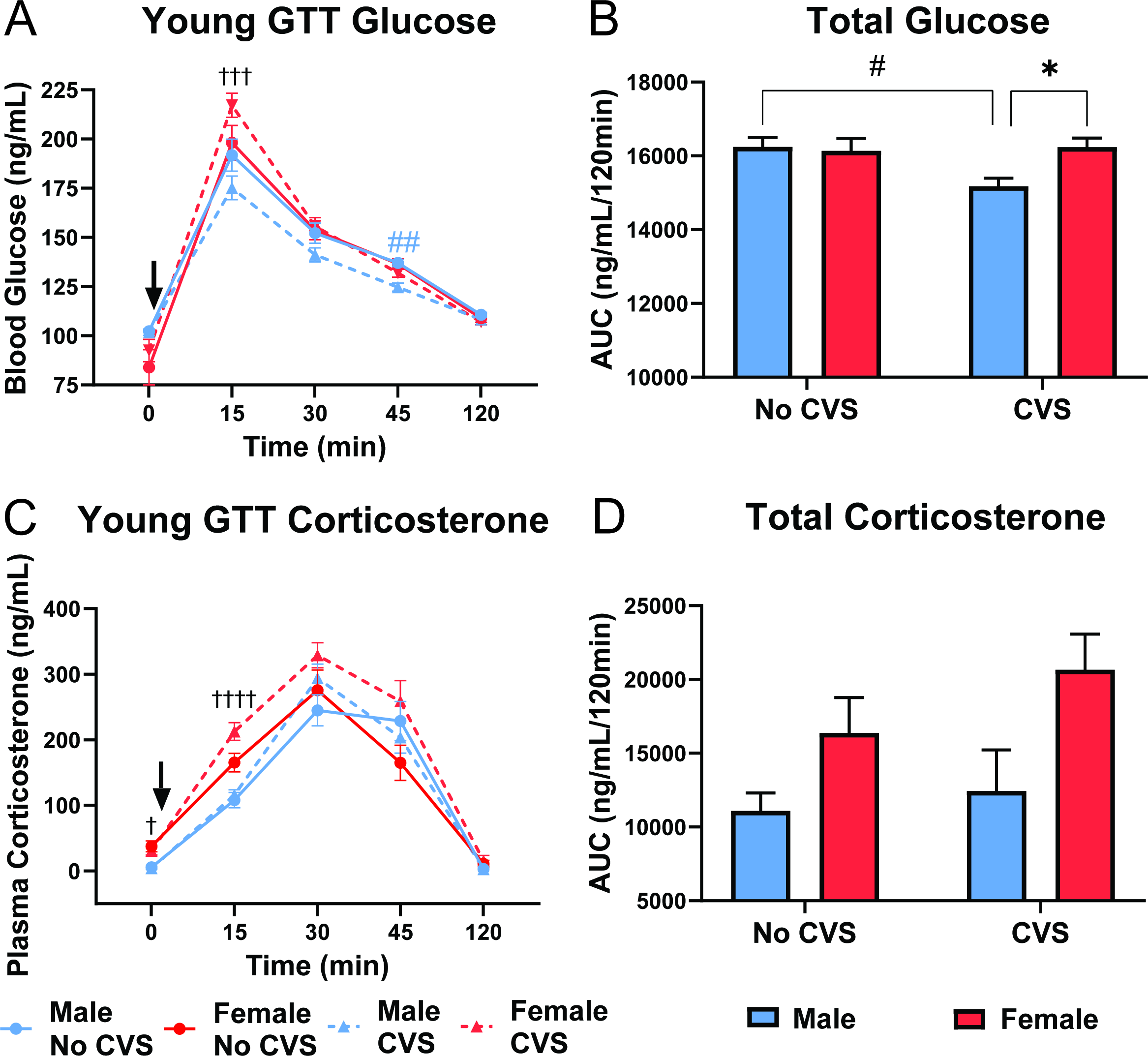
Young GTT. Basal blood samples (0 min) were taken before glucose injection (1.5 g/kg, i.p.). Blood was taken at 15, 30, 45, and 120 min post injection in No CVS (n = 24/sex) and CVS (n = 36/sex) rats. At each time point, blood glucose was measured (**A**) and total blood glucose was calculated from the AUC (**B**). Plasma corticosterone was also measured (**C**) and cumulative corticosterone determined from the AUC (**D**). Data are expressed as mean ± SEM. ↓ represents glucose bolus, * represents sex differences within stress condition, ^#^ represents CVS effects within sex (indicated by color for time-dependent measures), ^†^ represents sex differences within CVS over time. *,^#^,^†^ p < 0.05, ^##^ p < 0.01, ^†††^ p< 0.001, and ^††††^ p < 0.0001.

In response to glycemic challenge, corticosterone (**Fig. 4C**) had sex [F(1, 115) = 4.012, p = 0.0475], CVS [F(1, 115) = 6.002, p = 0.0158], and time [F(2.292, 199.4) = 165.7, p < 0.0001] effects with a CVS x time interaction [F(4, 348) = 3.108, p = 0.0156]. Post-hoc analysis revealed that, within CVS, females had higher (p < 0.05) fasting basal corticosterone than males; additionally, female CVS rats had higher corticosterone (p < 0.0001) at 15 min than CVS males. Analysis of total corticosterone (**Fig. 4D**) indicated a sex effect [F(1, 35) = 5.687, p = 0.0226]; however, females were not significantly different from males within either stress condition. Altogether, data from the GTT indicate that CVS improves male glucose clearance without affecting glucocorticoid responses. In contrast, CVS exposed females have impaired glucose tolerance and elevated glucocorticoid reactivity compared to their male counterparts.

### Aged body weight, corticosterone, and somatic measures

With aging and increased body weight, the sex difference in body weight persisted (**Fig. 5A**). ANOVA indicated effects of sex [F(1,115) = 1115, p < 0.0001] with males weighing more (p < 0.0001) than females in both stress conditions. While basal corticosterone (**Fig. 5B**) robustly increased in all groups, ANOVA found sex effects [F(1,115) = 26.04, p < 0.0001] where females had higher (p <0.01) resting corticosterone than males regardless of stress exposure. Somatic measures were determined at the conclusion of the study to investigate potential effects of sex and/or stress on adrenal weight, as well as body composition in terms of visceral mWAT and subcutaneous iWAT. Body weight-corrected adrenal indices (**Fig. 5C**) had a sex effect [F(1, 113) = 251.6, p < 0.0001] where females had greater (p <0.001) relative adrenal mass. There were no effects on body weight-corrected mWAT (**Fig. 5D**); in contrast, iWAT (**Fig. 5E**) had sex effects [F(1, 112) = 93.06, p < 0.0001] where females had less (p < 0.001) subcutaneous adiposity. In all, there were no effects of CVS on basal corticosterone or somatic measures. However, sex effects were present where females had lower body weight and less subcutaneous fat while their adrenal glands were larger and secreted more corticosterone.

**Figure 5:**
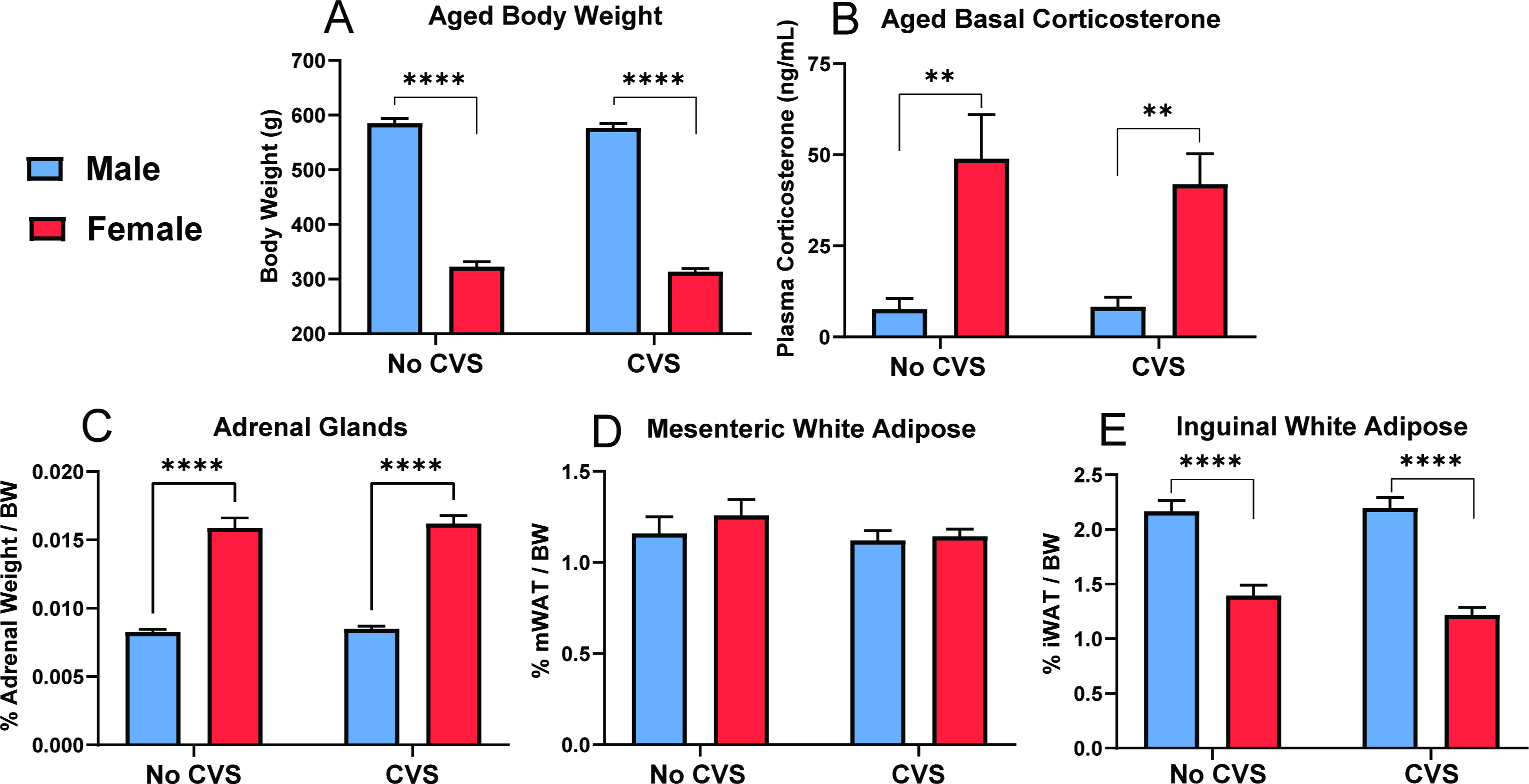
Aged basal and somatic measures. Following 13 months of aging, body weight (**A**) and basal corticosterone (**B**) were measured. Following euthanasia, body weight-corrected weights of adrenal glands (**C**), mesenteric white adipose (**D**), and inguinal white adipose (**E**) were determined. No CVS: n = 24/sex and CVS: n = 36/sex. Data are expressed as mean ± SEM. * represents sex differences within stress condition. ** p < 0.01 and **** p < 0.0001.

### Aged FST

In contrast to young behavior, females were more active than males in the FST at 15 months. Immobility (**Fig. 6A**) showed sex effects [F(1, 115) = 36.98, p < 0.0001] with males more immobile (p <0.001) than females in both stress conditions. Swimming (**Fig. 6B**) also had a sex effect [F(1, 115) = 15.81, p < 0.001]. While there was no significant difference in No CVS groups (p = 0.085), CVS females swam more (p < 0.01) than CVS males.

**Figure 6:**
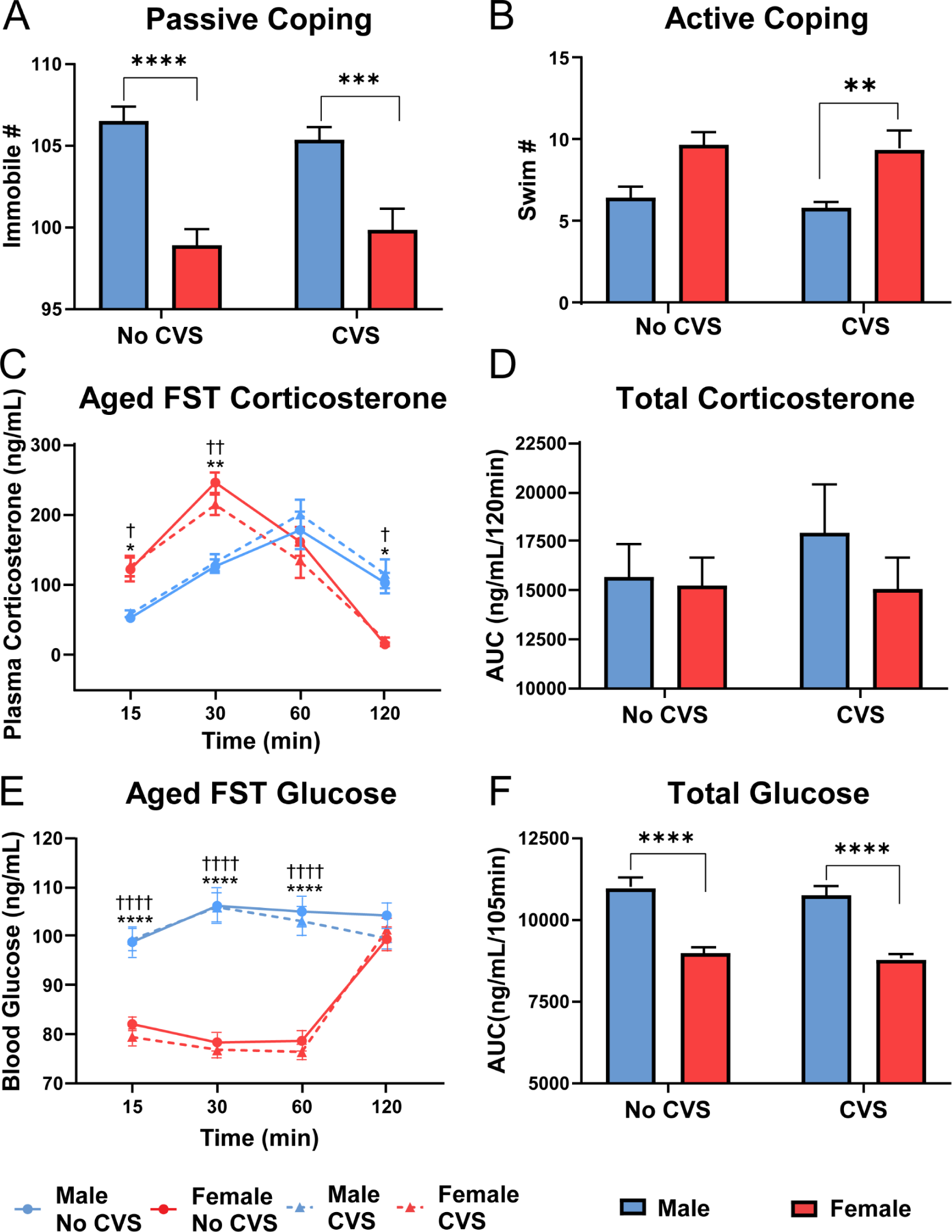
Aged FST. During FST, behavioral coping was assessed as passive (**A**) or active (**B**) in No CVS (n = 24/sex) and CVS (n = 36/sex) rats. Blood was taken at 15, 30, 60, and 120 min after the initiation of the 10-min FST. Plasma corticosterone was measured (**C**) and the total corticosterone response was calculated from the AUC (**D**). Blood glucose was also measured (**E**) and total blood glucose calculated (**F**). Data are expressed as mean ± SEM. * represents sex differences within stress condition and ^†^ represents sex differences within CVS over time. *,^†^ p < 0.05, **,^††^ p < 0.01, *** p< 0.001, and ****,^††††^ p < 0.0001.

Corticosterone responses to FST (**Fig. 6C**) had time [F(2.52, 246.4) = 114.7, p < 0.0001] and sex x time [F(4, 391) = 38.08, p < 0.0001] effects. Females in both stress groups generated more rapid corticosterone responses with elevated levels at 15 (p < 0.05) and 30 (p < 0.01) min. However, delayed male recovery from the stressor led to females having lower (p < 0.05) corticosterone at 120 min. Given that females had greater corticosterone responses immediately after the stressor and that males showed delayed responses, AUC analysis (**Fig. 6D**) indicated similar total corticosterone exposure in all groups.

Glucose responses to FST in aged animals (**Fig. 6E**) had sex [F(1, 116) = 77.58, p < 0.0001] and time effects [F(2.022, 233.1) = 44.90, p < 0.0001] with a sex x time [F(3, 346) = 61.63, p < 0.0001] interaction. As seen in young animals, males had higher glucose (p < 0.001) at 15-60 min in both stress groups. This was reflected in the AUC (**Fig. 6F**) where main effects of sex [F(1, 114) = 72.41, p < 0.0001] where present in both stress conditions. In summary, females had less immobility in the FST with CVS females more actively coping. This was coupled with an age-related shift in the temporal dynamics of male and female glucocorticoid responses where females had more rapid stress responses.

### Aged GTT

After aging, the GTT was repeated and analysis of glucose clearance (**Fig. 7A**) found time effects [F(1.66, 192.4) = 407.9, p < 0.0001] with no significant group differences. There were also no significant differences in total glucose (**Fig. 7B**).

**Figure 7:**
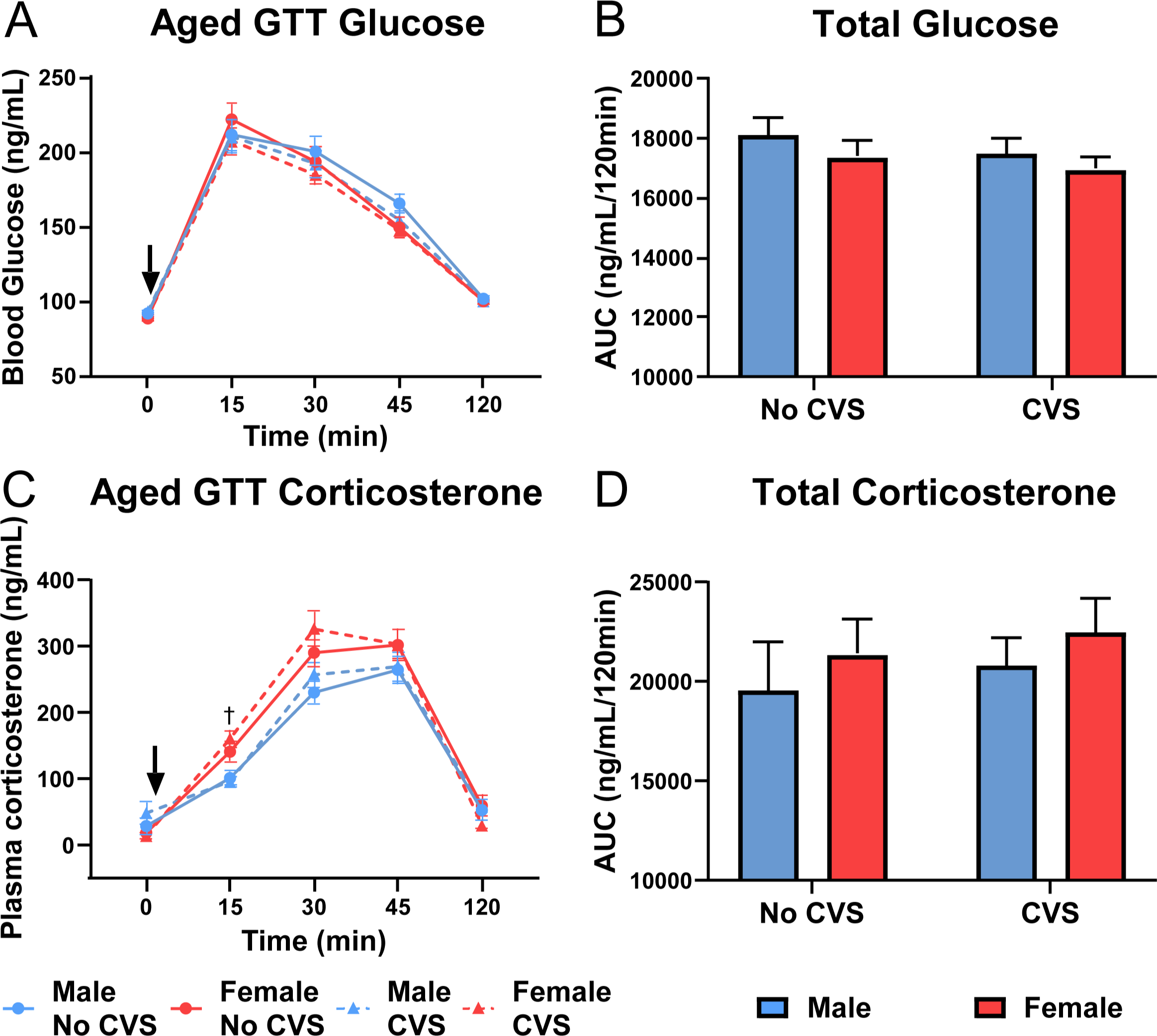
Aged GTT. Basal blood samples (0 min) were taken before glucose injection (1.5 g/kg, i.p.). Blood was taken at 15, 30, 45, and 120 min post injection in No CVS (n = 24/sex) and CVS (n = 36/sex) rats. At each time point, blood glucose was measured (**A**) and total blood glucose was calculated from the AUC (**B**). Plasma corticosterone was also measured (**C**) and cumulative corticosterone determined from the AUC (**D**). Data are expressed as mean ± SEM. ↓ represents glucose bolus, represents sex difference within CVS (p < 0.05).

Examination of glucocorticoid responses to glycemic challenge (**Fig. 7C**) indicated stress [F(1, 115) = 5.184, p = 0.0246] and time [F(2.83, 251.2) = 253.6, p < 0.0001] effects with stress x time [F(4, 355) = 6.055, p = 0.0001] interactions. Specifically, females exposed to CVS had elevated corticosterone at 15 min compared to CVS males. Although, there were no significant differences in AUC (**Fig. 7D**). Thus, there were no group differences in aged glucose tolerance, but chronically stressed females maintained greater glucocorticoid reactivity.

### Estrous cycle

Estrous cycle was determined for females following each acute stress test (**Table 1**). Given the single cohort design of the study, cycling was random and not staged.

**Table 1:** Percentage of animals in each estrous cycle phase during acute tests.

|  | Phase | Young FST | Young GTT | Aged FST | Aged GTT |
| --- | --- | --- | --- | --- | --- |
| **No CVS** | Proestrus | 16.66 | 33.33 | 20.83 | 8.33 |
|  | Estrus | 75 | 45.83 | 62.5 | 75 |
|  | Metestrus | 4.16 | 20.83 | 8.33 | 8.33 |
|  | Diestrus | 4.16 | 0 | 8.33 | 8.33 |
| **CVS** | Proestrus | 16.66 | 13.88 | 27.77 | 22.22 |
|  | Estrus | 77.77 | 69.44 | 44.44 | 38.88 |
|  | Metestrus | 5.55 | 8.33 | 22.22 | 16.66 |
|  | Diestrus | 0 | 8.33 | 5.55 | 22.22 |

However, lifelong cohabitation increased synchronization. For instance, 46 of 60 females were in estrus for young FST. With aging, there was more spread across cycles, especially in the CVS group. As most assessments included rats in all phases, cycle phase was included in correlational analyses to investigate potential effects on behavior and endocrine outcomes. Surprisingly, estrous cycle phase did not correlate with coping behavior or endocrine responses; although, cycle phase during each assessment correlated with phase during other assessments.

### Regression analysis: No CVS males

Pearson’s correlations were conducted to examine potential associations between behavioral and endocrine data with an emphasis on how early life measures predict later life outcomes. Endocrine responses during FST and GTT were represented by the total AUC for each assessment. All correlations are visible in figure 8 with a subset discussed below.

**Figure 8:**
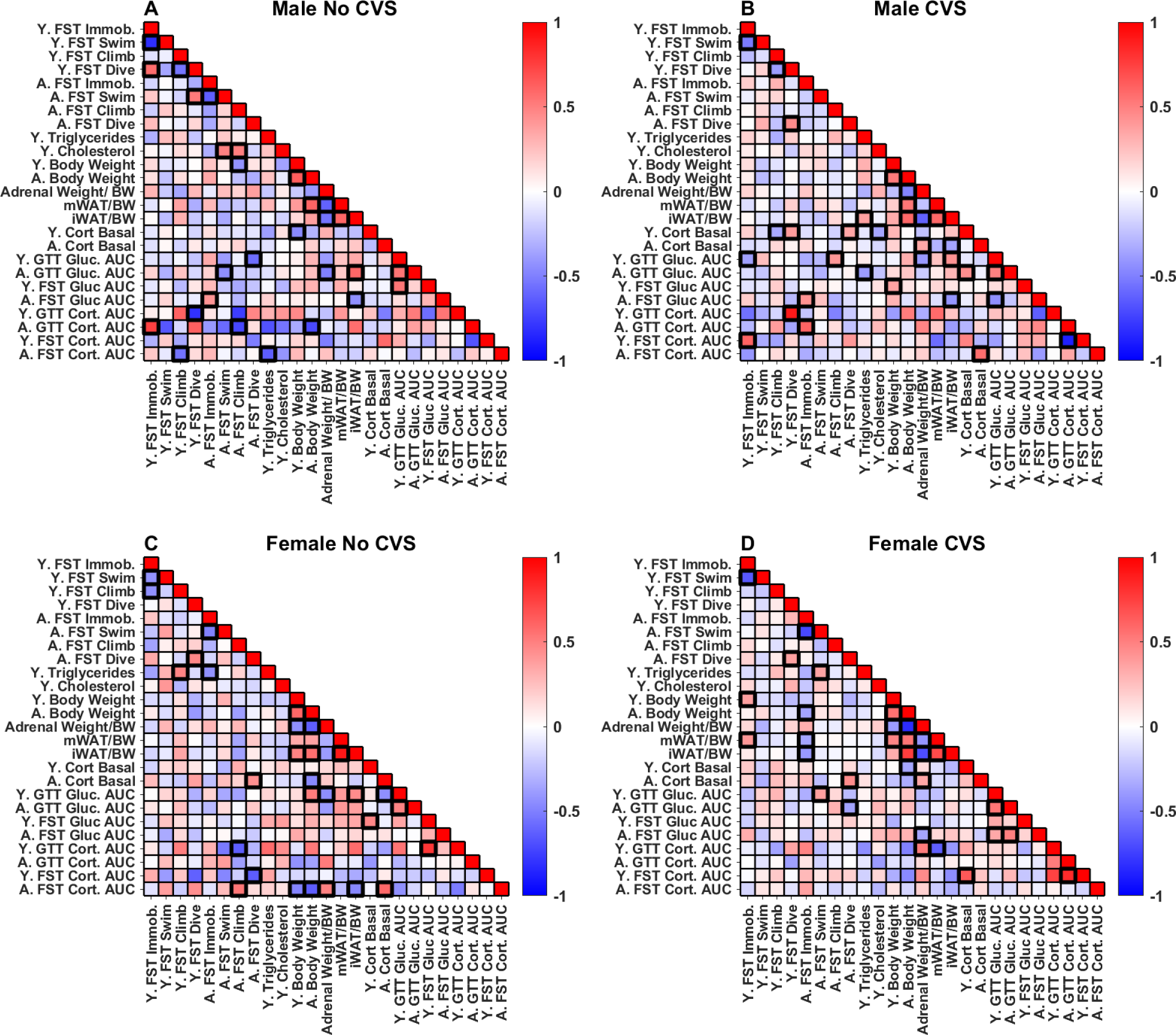
Regression analysis. Pearson correlations were run within each group [No CVS (n = 24/sex) and CVS (n = 36/sex)] for all measures taken throughout the study. Significant associations (p < 0.05) are bolded.

In No CVS males (**Fig. 8A**), there were a limited number of associations within data collected in young animals. Glucose responses to FST and GTT were positively correlated (r = 0.536, p = 0.008). Additionally, there were negative correlations between body weight and basal corticosterone (r = -0.463, p = 0.023), as well as diving behavior and corticosterone responses to hyperglycemia (r = -0.475, p = 0.028).

Within data collected in aged animals, there was a positive correlation between immobility and glucose in the FST (r = 0.415, p = 0.048). Furthermore, swimming and climbing in the FST negatively correlated with GTT glucose (r = -0.437, p = 0.036) and corticosterone (r = -0.72, p = 0.044), respectively, suggesting that active coping behaviors associate with reduced responses to glycemic challenge in aged males.

Multiple correlates of somatic measures were present, including negative associations between adrenal weight and glucose during GTT (r = -0.498, p = 0.013), iWAT (r = -0.547, p = 0.007), and mWAT (r = -0.615, p = 0.001). Subcutaneous iWAT also had a positive relationship with glucose in the GTT (r = 0.594, p = 0.002) and a negative relationship with FST-evoked glucose (r = -0.437, p = 0.036).

In terms of young data that predict aged outcomes, young immobility positively correlated with aged corticosterone responses to glycemic challenge (r = 0.74, p = 0.034). Also, active climbing in the young FST negatively correlated with aged FST corticosterone (r = -0.548, p = 0.028). Together, these associations indicate that young coping behaviors predict aged glucocorticoid responses to systemic and psychological stressors. Interestingly, young basal cholesterol predicted increased swimming (r = 0.495, p = 0.019) and climbing (r = 0.502, p = 0.017) in the aged FST, while young basal triglycerides negatively correlated with aged FST corticosterone (r = -0.636, p = 0.011). Although young and aged glucose tolerance were correlated (r = 0.546, p = 0.006), there were no associations at any age between glucose and corticosterone, suggesting that the acute glucose responses assessed relate more to sympathetic than glucocorticoid control.

### Regression analysis: CVS males

In males exposed to chronic stress (**Fig. 8B**), numerous associations emerged that were not present in unstressed males. Among data collected in young animals, immobility in the FST positively correlated with corticosterone responses to FST (r = 0.612, p = 0.019). In contrast, immobility negatively correlated with glucose in the GTT (r = -0.372, p = 0.025). Basal corticosterone in young CVS males correlated with multiple outcomes including a positive association with diving behavior (r = 0.392, p = 0.018) and negative associations with climbing (r = -0.337, p = 0.044) and cholesterol (r = -0.354, p = 0.034).

Data collected from aged CVS males indicated that immobility in the FST positively correlated with glucose responses to FST (r = 0.428, p = 0.009) and corticosterone responses to glycemic challenge (r = 0.611, p = 0.026). Additionally, aged basal corticosterone related to adrenal weight (r = 0.354, p = 0.046) and FST corticosterone responses (r = 0.567, p = 0.018), while negatively correlating with iWAT (r = -0.359, p = 0.037). Overall, both young and aged CVS males had positive associations between passive coping and glucocorticoid responses to stressors.

As far as data collected in young animals immediately following CVS that predicted outcomes in aged rats, glucose in the young GTT correlated positively with aged GTT (r = 0.495, p = 0.002) and climbing behavior (r = 0.407, p = 0.013) but negatively with FST glucose (r = -0.438, p = 0.007). Young triglycerides predicted iWAT (r = 0.371, p = 0.018) and negatively associated with glucose in the aged GTT (r = -0.386, p = 0.019). Additionally, young basal corticosterone during CVS predicted GTT glucose (r = 0.448, p = 0.006) and diving behavior in aged rats (r = 0.349, p = 0.037). While unstressed males did not have associations at any age between glucocorticoid responses to psychological and physiological stressors, CVS males had a negative correlation between young FST corticosterone responses and aged corticosterone responses to hyperglycemia (r = -0.873, p = 0.023).

### Regression analysis: No CVS females

In young female controls (**Fig. 8C**), triglycerides positively correlated with FST climbing (r = 0.48, p = 0.021). Additionally, basal corticosterone positively associated with glucose mobilization during the FST (r = 0.466, p = 0.022), which associated with corticosterone responses to GTT (r = 0.79, p = 0.004). Together, these findings indicate that, in contrast to males, female endocrine responses to swim and glycemic stressors are positively related.

In aged No CVS females there were numerous relationships between glucocorticoids and coping behaviors. Specifically, aged basal corticosterone positively correlated with diving (r = 0.436, p = 0.038) and FST corticosterone responses (r = 0.593, p = 0.007). Further, aged FST glucocorticoid responses correlated with climbing behaviors (r = 0.538, p = 0.018). The aged FST glucocorticoid response also related positively to adrenal weight (r = 0.495, p = 0.031) and negatively to body weight (r = -0.635, p = 0.003) and iWAT (r = -0.473, p = 0.041). Ultimately, these correlations suggest that adrenal glucocorticoid release is related to active coping in aged females.

Young female data that predicted aged outcomes include corticosterone responses to both glycemic and psychogenic stressors negatively correlating with active coping behaviors after aging. Specifically, young corticosterone responses to hyperglycemia were negatively correlated with aged climbing behavior (r = -0.630, p = 0.037), while corticosterone responses to forced swim were negatively correlated with diving behavior (r = -0.636, p = 0.047). In addition, young triglycerides were negatively correlated with immobility after aging (r = -0.431, p = 0.040). Collectively, these correlations suggest that the relationship between HPA axis activity and coping behavior changes with age. That is, aged females show a positive relationship between active coping and corticosterone responses; in contrast, increased glucocorticoid responses immediately after CVS negatively associate with active coping later in life.

### Regression analysis: CVS females

In young chronically-stresses females (**Fig. 8D**), there were no associations between behavioral and endocrine measures. In fact, the only correlations among data collected following CVS were positive relationships between basal corticosterone and corticosterone responses to FST (r = 0.652, p = 0.005) and immobility in the FST relating to body weight (r = 0.37, p = 0.026). When compared to No CVS females, these data suggest that prolonged stress exposure may disrupt the female-specific relationships between endocrine responses to psychological and physiological stressors.

With aging, more relationships arose including passive coping negatively relating to body weight (r = -0.36, p = 0.031) and adiposity (mWAT: r = -0.355, p = 0.034, iWAT: r = -0.403, p = 0.017). Additionally, glucose in the FST and GTT correlated (r = 0.502, p = 0.002), although there was no relationship between glucocorticoid responses in these tests. Moreover, diving during the FST positively correlated with basal corticosterone (r = 0.453, p = 0.006) and negatively correlated with GTT glucose (r = -0.362, p = 0.03).

Young measures that predicted aged responses included glucose during the young GTT correlating with aged active coping (swim: r = 0.375, p = 0.024) and glucose during aged FST (r = 0.342, p = 0.044) and GTT (r = 0.535, p = 0.0007). Swimming during the aged FST was also positively associated with young triglycerides (r = 0.361, p = 0.03). In contrast to both No CVS females and chronically-stressed males, CVS females had correlates of aged visceral adiposity. In fact, young immobility correlated positively (r = 0.41, p = 0.013) and young GTT corticosterone negatively (r = -0.628, p = 0.009) with mWAT. Collectively, these findings indicate that chronic stress has sex-specific effects on the relationships between passive coping, glucose tolerance, and visceral adiposity over the lifespan.

## Discussion

Broadly, this study sought to examine the behavioral and metabolic consequences of adolescent chronic stress in male and female rats across the lifespan. Compared to males immediately after CVS exposure, females exhibited elevations in basal corticosterone, passive coping behavior, corticosterone responses to behavioral stress, and hyperglycemia-evoked corticosterone, in addition to impaired glucose tolerance. While there were prominent sex by age interactions, enduring sex differences in the effects of chronic stress were present in coping behavior and corticosterone responses to glycemic challenge. Interestingly, age impacted sex differences in coping behavior, as young females had more passive coping while males were more passive after aging. In fact, regression analysis indicated that young and aged coping behaviors had little relation in any group, suggesting this behavior may represent more of a state than trait variable. In contrast, glucose tolerance was a stable trait across the lifespan in all groups. Collectively, the results indicate that adolescent chronic stress exposure has sex-specific impacts on behavior and metabolic health, partially contributing to differential outcomes later in life.

Adolescent chronic stress exposure leads to a number of physiological changes (Cotella et al., 2020; Foilb et al., 2011; Jankord et al., 2011; McEwen, 2009; Romeo et al., 2007, 2004; Vargas et al., 2016; Wulsin et al., 2016), which held true in this study. Particularly, chronically stressed females had increased glucocorticoid responses across the life span which may indicate a broad sensitization of the HPA axis.

Additionally, chronically stressed young females had impaired glucose clearance. Taken together, this phenotype may represent impaired metabolic capacity and/or overall dysregulation of endocrine homeostasis. Further, regressive analysis indicated that, in chronically stressed females, metabolic indicators including triglycerides and blood glucose were positively correlated with active coping behaviors after aging.

Although numerous studies have reported increased passive coping in females at varying time points after adolescent chronic stress, effects on glucocorticoid responses have been mixed (Cotella et al., 2020; Smith et al., 2018; Wulsin et al., 2016). Specifically, adolescent chronic stress in female rats has been found to produce both hypoactivity (Wulsin et al., 2016) and sensitization (Cotella et al., 2020) of the HPA axis. The present results found female HPA axis sensitization to both behavioral and glycemic stressors immediately after chronic stress that persisted, in part, into late adulthood. Although timing and age differences in stress exposure/data collection may contribute to divergent results, the current findings of glucose intolerance combined with widespread female HPA axis sensitization suggest greater overall endocrine and autonomic reactivity in stressed females.

Interestingly, chronic stress exposure improved glucose tolerance in young males. This associated negatively with young FST immobility and positively with body weight-corrected subcutaneous adiposity at the end of the study, correlations that did not exist in No CVS males. It remains to be determined whether relative differences in adiposity following CVS play a role in the increase rated of glucose clearance.

Androgen effects are also possible as testosterone improves glucose tolerance in men (Navarro et al., 2015), and increases active coping in male rats (Frye and Wawrzycki, 2003).

In both young male and female rats, CVS impacted behavioral and endocrine measures, including elevated glucocorticoid responses to behavioral stress. However, sex differences in the effects of aging more prominently impacted long-term outcomes than CVS. Regardless of stress status, males and females differed in basal glucocorticoid tone, subcutaneous adiposity, coping behavior, and stress-induced glucose mobilization. In fact, the increased immobility observed in young females was contrasted by an age-related increase in male immobility. This may reflect androgen decline in males and/or stress resilience in aged females (Hodes and Epperson, 2019). Further, the central processing of stress would be expected to contribute to sex-and stress-specific effects. In fact, 6 weeks after adolescent CVS, FST-induced Fos expression shows sex differences in chronic stress responses. Here, male rats have decreased neural activation in the medial and basolateral amygdala, while females do not (Cotella et al., 2020).

Although glucose tolerance is a well-accepted measure of metabolic function (Hahn et al., 2011), numerous other hormones are critical, most notably insulin. Given the design of sampling blood 5 times each in 2 experiments during the same week, we were limited in the volume we were able to collect. That combined with the demands of processing 2,400 total blood samples across the study, meant sample measurements were prioritized to prominent indicators of both stress reactivity and metabolic regulation. More targeted analysis of insulin in future studies would be expected to better illuminate sex differences in glucose handling, as well as potential effects on insulin sensitivity. Another design consideration relates to allowing animals to age to 15 months, which should near reproductive senescence (Brinton, 2012). This was chosen as a relative translational equivalent of late middle-aged. Expanding the age range beyond 18 months would ensure reproductive senescence and may yield differences due to the reduced levels of gonadal hormones at advanced age. Despite the limitations, this study provides novel insights into the longitudinal effects of adolescent chronic stress and a basis for understanding sex-specific pathologies.

Epidemiologic data indicate that males and females develop metabolic syndrome at similar rates (Ford et al., 2002). However, metabolic syndrome is a predisposing factor to cardiovascular disease (Saklayen, 2018), which is an increasingly prevalent cause of death in women after midlife (Rosamond et al., 2007; Towfighi et al., 2009).

Our data suggest that prolonged stress exposure may potentiate physiological and behavioral changes associated with the development of metabolic syndrome and subsequent disease sequalae. Further investigation may indicate that sex-specific predictive correlations support biomarker development to predict or intervene in future pathology. Given the role of chronic stress in psychiatric and metabolic disorders (Chandola et al., 2006; Fiksdal et al., 2019; Picard et al., 2014; Yusef et al., 2004), mitigating the consequences has the potential to improve health and quality of life.

## Acknowledgements

The authors are grateful for the colleagues that made the scale of these experiments possible. We would like to thank (in alphabetical order) Jody Caldwell, Fernanda Correa, Evelin Cotella, Nathan Evanson, Sarah Fourman, Amy Packard, Benjamin Packard, Brittany Smith, and Aynara Wulsin for assistance with blood sampling and tissue collection. Additionally, Sebastian Pace aided in data management and Thomas Dearing assisted with regression analysis.

## Funding

This work was supported by NIH grants HL122454 and HL150559 to B. Myers and a Pilot Translation Research Program grant to B. Myers and L. Wulsin from the University of Cincinnati Department of Psychiatry and Behavioral Neuroscience. Data generated by the University of Cincinnati Mouse Metabolic Phenotyping Core were supported by NIH grant DK059630. C. Dearing was supported by NIH institutional training grant GM136628 (PI: VandeWoude).

## Notes

### Competing Interest Statement

The authors have declared no competing interest.

